# Correcting palindromes in long reads after whole-genome amplification

**DOI:** 10.1101/173872

**Authors:** Sven Warris, Elio Schijlen, Henri van de Geest, Rahulsimham Vegesna, Thamara Hesselink, Bas te Lintel Hekkert, Gabino Sanchez Perez, Paul Medvedev, Kateryna D. Makova, Dick de Ridder

**Affiliations:** Applied Bioinformatics, Wageningen University and Research, Wageningen, the Netherlands; Bioinformatics Group, Wageningen University and Research, Wageningen, the Netherlands; Bioinformatics and Genomics Graduate Program, Pennsylvania State University, University Park, Pennsylvania 16802, USA; Computation, Bioinformatics, Statistics Graduate Training Program, Pennsylvania State University, University Park, Pennsylvania 16802, USA; The Genome Sciences Institute of the Huck Institutes of the Life Sciences, Pennsylvania State University, University Park, Pennsylvania 16802, USA; Department of Computer Science and Engineering, Pennsylvania State University, University Park, Pennsylvania 16802, USA; Department of Biochemistry and Molecular Biology, Pennsylvania State University, University Park, Pennsylvania 16802, USA; The Center for Medical Genomics, Pennsylvania State University, University Park, Pennsylvania 16802, USA; Department of Biology, Pennsylvania State University, University Park, Pennsylvania 16802, USA

## Abstract

Next-generation sequencing requires sufficient DNA to be available. If limited, whole-genome amplification is applied to generate additional amounts of DNA. Such amplification often results in many chimeric DNA fragments, in particular artificial palindromic sequences, which limit the usefulness of long reads from technologies such as PacBio and Oxford Nanopore. Here, we present Pacasus, a tool for correcting such errors in long reads. We demonstrate on two real-world datasets that it markedly improves subsequent read mapping and *de novo* assembly, yielding results similar to these that would be obtained with non-amplified DNA. With Pacasus long-read technologies become readily available for sequencing targets with very small amounts of DNA, such as single cells or even single chromosomes.

## Introduction

Modern sequencers require sufficient material to work with: the Illumina and Pacific Bioscience (PacBio) platforms prescribe at least three micrograms, but recommend at least five (Buermans and den Dunnen 2014) micrograms. Long-read sequencing technologies such as those offered by PacBio and Oxford Nanopore Technology (ONT) additionally require high molecular weight (HMW) DNA as a starting material, i.e. material in which individual DNA stretches are long. In many biological settings, obtaining sufficient amounts of DNA of the required quality and length is problematic, such as in studies on single cells (Shapiro et al. 2013; Gawad et al. 2016) or single selected chromosomes (Tomaszkiewicz et al. 2016). To overcome this limitation DNA is amplified, starting from as little as picograms, in a process called whole-genome amplification (WGA) (Czyz et al. 2015).

A major issue with the WGA process is that it introduces specific chimeric fragments (Lasken and Stockwell 2007; Sabina and Leamon 2015) consisting of one or more inverted repeats (Figure 1), so-called palindromes. This effect is partially alleviated by de-branching, however, chimeric fragments still remain (Zhang et al. 2006). In Illumina paired-end (PE) sequencing these fragments are then sheared into small sub-fragments before library preparation, which reduces the effect on subsequent analyses of the palindromic fragments as they will occur in only few reads. In other approaches to sequencing, however, the full fragments are used. For Illumina mate-pair (MP) sequencing, long palindromic fragments will result in pairs with incorrect directions and unpredictable insert sizes. As a result short read MP libraries based on WGA are problematic for read mapping and *de novo* assembly.

**Figure 1:**
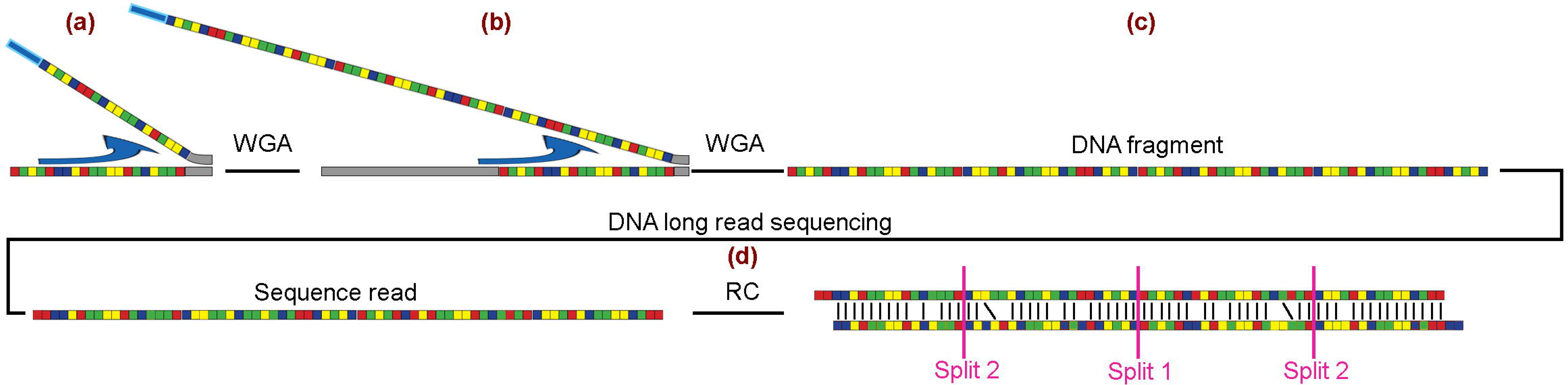
The introduction of palindromes by whole-genome amplification (WGA) and correction of these sequences with Pacasus. The colored squares in this figure indicate the four different nucleotides. Palindromes are introduced when during WGA the DNA-polymerase continues with elongation (indicated by the arrow) along an already created WGA product (a), generating a palindrome. In this example this incorrect elongation occurs several times (b), resulting in a DNA fragment containing four copies of the original fragment (c), which is sequenced. Pacasus detects the palindrome sequence by aligning the read’s reverse complement to itself (d) and splits the read in two smaller reads at the center of the alignment (split 1). This process is repeated and splits the two resulting reads again (split 2), yielding four separate, ‘clean’, reads. The full set of reads, corrected and left intact, is then used in, for example, read mapping or *de novo* assembly.

Tools specifically aiming to detect and correct chimeric reads have been proposed (e.g. uchime (Edgar et al. 2011)) and work well for paired-end and single-end short-read technologies.

For long reads however, the palindromic nature of the sequence hinders read mapping and renders *de novo* assembly highly problematic. Due to the high base-calling error rate of the long read technologies (11-38%) (Rhoads and Au 2015; Jain et al. 2016), finding and correcting these palindromic constructs in long reads cannot be done by exact string matching. Algorithms for improving long-read quality in general are available: Proovread (Hackl et al. 2014), PBcR (Koren et al. 2012) and ECTools (Lee et al. 2014) use either Illumina HiSeq reads or assembled contigs based on HiSeq data to increase the quality of base calling. While Proovread can also detect chimeric fragments, it specifically aims at detecting PCR artifacts joining fragments originating from different regions in the genome. This is done by mapping HiSeq reads, assumed to be available, which do not have these chimeras. The location of the chimera in the long read is then detected by finding discrepancies in the short-read mappings. This approach is unfit for solving the chimeras occurring due to WGA: the HiSeq reads are based on the same fragments as the long reads and will therefore contain the same nucleotide sequence information. As a consequence of this lack of suitable methods to correct these chimeras, the use of WGA with long-read technologies was usually not advised (Lasken and Stockwell 2007), which precludes the application of long read technology to answer essential biological questions at the single-chromosome or single-cell level.

Here, we introduce a new method, *Pacasus*, for correcting palindromic, long, error-rich reads without the loss of nucleotide information. The method relies on a Smith-Waterman alignment implementation called pyPaSWAS (Warris et al. 2015; Warris and Timal 2016), which supports fast processing on multicore CPUs, GPUs and Xeon Phis to detect palindromes and iteratively corrects them by splitting up reads. We demonstrate its performance on PacBio sequencing data of *Arabidopsis thaliana* as well as on flow-sorted gorilla Y chromosome data.

The gorilla Y chromosome was selected because primate Y chromosomes are relatively short and contain many repeats, rendering them difficult to sequence and assemble. Even in one of the most complete assemblies, that of *Homo sapiens*, more than half of the sequence of Y is still unknown (Human Genome Sequencing Consortium International 2004). To obtain a higher read coverage of the gorilla Y chromosome, a recent paper (Tomaszkiewicz et al. 2016) used flow-sorting and WGA. PacBio long reads, Illumina HiSeq PE and MP-libraries, transcriptome data and PCR sequences were used by the authors as well (Bioproject PRJNA293447). The RecoverY tool presented in same paper was designed to identify short reads originating from the Y chromosome. Based on these data, the authors created a hybrid (PacBio + HiSeq) assembly, here labeled ‘GorY’. The authors also used HGAP (Chin et al. 2013) and MHAP (Berlin et al. 2015) to create PacBio-only assemblies, but these resulting assemblies were of suboptimal quality. In this manuscript, we used the PacBio data after WGA generated by Tomaszkiewicz and colleagues (Tomaszkiewicz et al. 2016) to show the benefits of correcting palindrome sequences in this data set with an improved quality of the PacBio-only *de novo* genome assembly.

## Results

### Pacasus corrects many palindromic sequences found in WGA data

To demonstrate the added value of Pacasus in the analysis of PacBio reads generated from WGA DNA samples, we applied it to several data sets of *Arabidopsis thaliana* and a data set of the gorilla Y chromosome (Tomaszkiewicz et al. 2016) (Table 6). Table 1 lists the original number of reads, the number of reads that were found to have chimeras and the number of clean reads after correcting the palindromes. In the Arabidopsis samples, 40-50% of reads contained at least one palindrome, with some reads containing up to 15 (Suppl. Fig. 1). This demonstrates the extent to which palindromes pose a problem in PacBio WGA data and illustrates that Pacasus effectively detects and corrects these.

**Table 1.** Effect of correcting palindromes on the number reads and average lengths of these reads.

| Sample | Before cleaning |  | Reads with detectable palindromes |  | After correcting |  |
| --- | --- | --- | --- | --- | --- | --- |
|  | Number of reads | Average length (b) | Number of reads | Number of reads (%) | Number of reads | Average length (b) |
| Ath-WGA1 | 462138 | 9326 | 221001 | 47.8 | 869826 | 4660 |
| Ath-WGA2 | 447364 | 8544 | 195263 | 43.6 | 769027 | 4721 |
| GorY-WGA | 3596236 | 5468 | 426188 | 11.8 | 4546488 | 4234 |

Pacasus decreases the average read length of the *A. thaliana* set by about 48%, i.e. preserving much of the long-range information. In the gorilla read set, 11.8% of the reads contain palindromes, which is less than are found in the *A. thaliana* sets. The average length of the gorilla reads before processing with Pacasus is 5468b, i.e. 61.2% of the average length in the total Ath-WGA data set (8934b); after correcting the palindromes this is increased to 90.3%: 4234b compared to 4689b.

### Correcting palindromes improves read mapping

Using the BLASR default settings and an additional filter of at least 80% nucleotide identity between read and reference, both the raw and clean read sets map well (Table 2). Palindromic reads map partially, leaving a (potentially large) proportion of the reads unmapped. This becomes clear when only mappings are considered where 80% and 95% of the complete read can be aligned: mapping efficiency for the raw read set drops from 99% to 44% and finally to 34%. For the clean reads, the mapping rates are 99%, 81% and 66% respectively, higher than for the noWGA read set (95%, 72% and 57%). Average coverages show similar effects. These mappings statistics indicate that the clean reads map more accurately and with higher read coverage than the raw reads do. The complete mapping reports are presented in Sup. reports 1-3.

**Table 2.** Statistics of read mappings with BLASR to the TAIR10 reference genome, calculated without filtering for a minimum read alignment length (‘-’) and after filtering for reads aligned with at least 80% or 95% nucleotide identity.

| Alignment filter | Reads mapped (%) |  |  | Avg. coverage |  |  | Avg. read length |  |  |
| --- | --- | --- | --- | --- | --- | --- | --- | --- | --- |
|  | - | 80% | 95% | - | 80% | 95% | - | 80% | 95% |
| Ath-WGA | 99 | 44 | 34 | 40.5 | 17.4 | 13.4 | 8987 | 5568 | 5397 |
| Ath-Clean | 99 | 81 | 66 | 55.1 | 48.1 | 41.2 | 4690 | 4532 | 4697 |
| Ath-Ctrl | 95 | 72 | 57 | 34.2 | 29.2 | 24.8 | 5799 | 5583 | 5852 |

To verify the palindromic nature of the reads, the locations of the clean reads were also investigated. If the raw reads indeed contain palindromic sequences, the parts of the clean reads should map to the same region in the genome (in contrast to chimeric reads, where the parts originate from different regions in the genome). To verify this, the longest distance between the mapping locations of each part of the corrected reads was calculated and related to the length of the original raw read. 96.5% of these mapping distances are within the read length of the original read, showing that most of the clean reads map to the same region in the genome and that the original raw reads indeed contain palindromes, not other types of chimeras.

### Assembly quality of corrected WGA reads approaches that of non-amplified data

To assess whether correcting palindromes also benefits assembly, we investigated two realistic scenarios using the *A. thaliana* data (Ath-WGA, Ath-Clean, Ath-Ctrl): PacBio-only assembly using Canu and a hybrid assembly, combining PacBio and Illumina HiSeq data, using DBG2OLC/Sparse. On the control data set Ath-Ctrl, this results in assemblies with overall good assembly statistics, with DBG2OLC yielding the best results (Table 3). Repeating the process with the original WGA data gives far worse results; in particular the DBG2OLC assembly has, for example, an N50 value about 26-fold smaller than the N50 value of the control data and the assembly covers about half (49.7%) of the reference genome.

**Table 3.** Statistics on the PacBio-only and hybrid assemblies of the various datasets. Note that the TAIR10 reference genome is 119.7Mb, with the full genome thought to be approximately 135Mb (Schmuths et al. 2004).

|  | PacBio-only (Canu) |  |  | Hybrid (DBG2OLC/Sparse) |  |  |
| --- | --- | --- | --- | --- | --- | --- |
| Read set | Ath-Ctrl<br>(C1) | Ath-WGA<br>(C2) | Ath-Clean<br>(C3) | Ath-Ctrl<br>(D1) | Ath-WGA<br>(D2) | Ath-Clean<br>(D3) |
| No. contigs | 852 | 2,128 | 1,015 | 476 | 4,818 | 1,753 |
| Ass. length (Mbp) | 115.6 | 116.8 | 123.9 | 110.9 | 108.9 | 131.0 |
| Longest contig (Kbp) | 1181 | 655 | 3402 | 5667 | 246 | 2239 |
| GC (%) | 36.0 | 36.2 | 36.12 | 35.97 | 36.57 | 36.21 |
| N50 (Kbp) | 293 | 73 | 302 | 823 | 32 | 278 |
| L50 | 117 | 479 | 109 | 33 | 951 | 113 |
| Covered (%) | 86.6 | 91.2 | 97.3 | 85.1 | 49.7 | 96.3 |
| Dupl. ratio | 1.09 | 1.07 | 1.06 | 1.08 | 1.34 | 1.13 |

Correcting the palindromic reads improves the hybrid assembly: although the N50 is lower than that of the Ath-Ctrl assembly, the assembly length and genome coverage are higher.

The Ath-Clean PacBio-only assembly is even better than the assembly based on the Ath-Ctrl data, with a higher N50 and genome coverage (Table 3). This is also reflected by the contig length distribution (Suppl. Fig. 2). Apparently the removal of conflicting information outweighs the loss of long-range information.

The hybrid assembly and the PacBio-only assembly based on Ath-Clean are longer in size than the TAIR10 reference genome (119.7Mb), being 123.9Mb and 131.0Mb respectively. The full genome is thought to be approximately 135Mb (Schmuths et al. 2004), so this additional sequence information could be new genomic data. No further testing has been done to verify this.

### *De novo* assembly based solely on long reads of flow-sorted chromosomes is now possible

After correcting the palindromes in the original gorilla PacBio reads (see section 3.1) we were able to create two PacBio-only assemblies: GorY-WGA based on the raw data set and GorY-Clean, based on the clean reads. The GorY-WGA assembly was added to the comparison to stay in line with the *Arabidopsis thaliana* analyses described in the previous section and also to verify that the increase in quality is not only due to a better performing software application. Figure 2 shows the length distributions of both the contigs and the scaffolds in the previously published GorY hybrid assembly and the contigs in the new GorY-Clean / Gory-WGA PacBio-only assemblies. The GorY scaffolds were created by using long-range information to connect the contigs (Tomaszkiewicz et al. 2016). The scaffolds contain no additional information, except sequence continuity. Gaps between contigs in the scaffolds are filled with Ns. The top-10 longest contigs of GorY-Clean are as long as the top-10 longest GorY scaffolds, showing that the new contigs already contain the same continuity except that the gaps are filled with sequence information. The scaffolded GorY assembly seems larger than the GorY-Clean one (Table 4, Suppl. Fig. 3). However, this is misleading as it contains 2.4 Mbp of Ns; the actual nucleotide content of the GorY assembly is 1.3 Mbp less than that of GorY-Clean. This is corroborated by further assembly statistics (Table 4). In terms of structure, the GorY-Clean assembly resembles the human Y chromosome assembly more than the original assembly (Suppl. Fig. 4). The GorY-WGA assembly is also of higher quality compared to the GorY contigs, but not as good as the GorY-Clean assembly. We attribute the quality increase of GorY-WGA compared to the GorY assembly to the use of Canu (Koren et al. 2017); the improvement of GorY-Clean over GorY-WGA is most likely due to correcting the palindromic reads with Pacasus.

**Figure 2:**
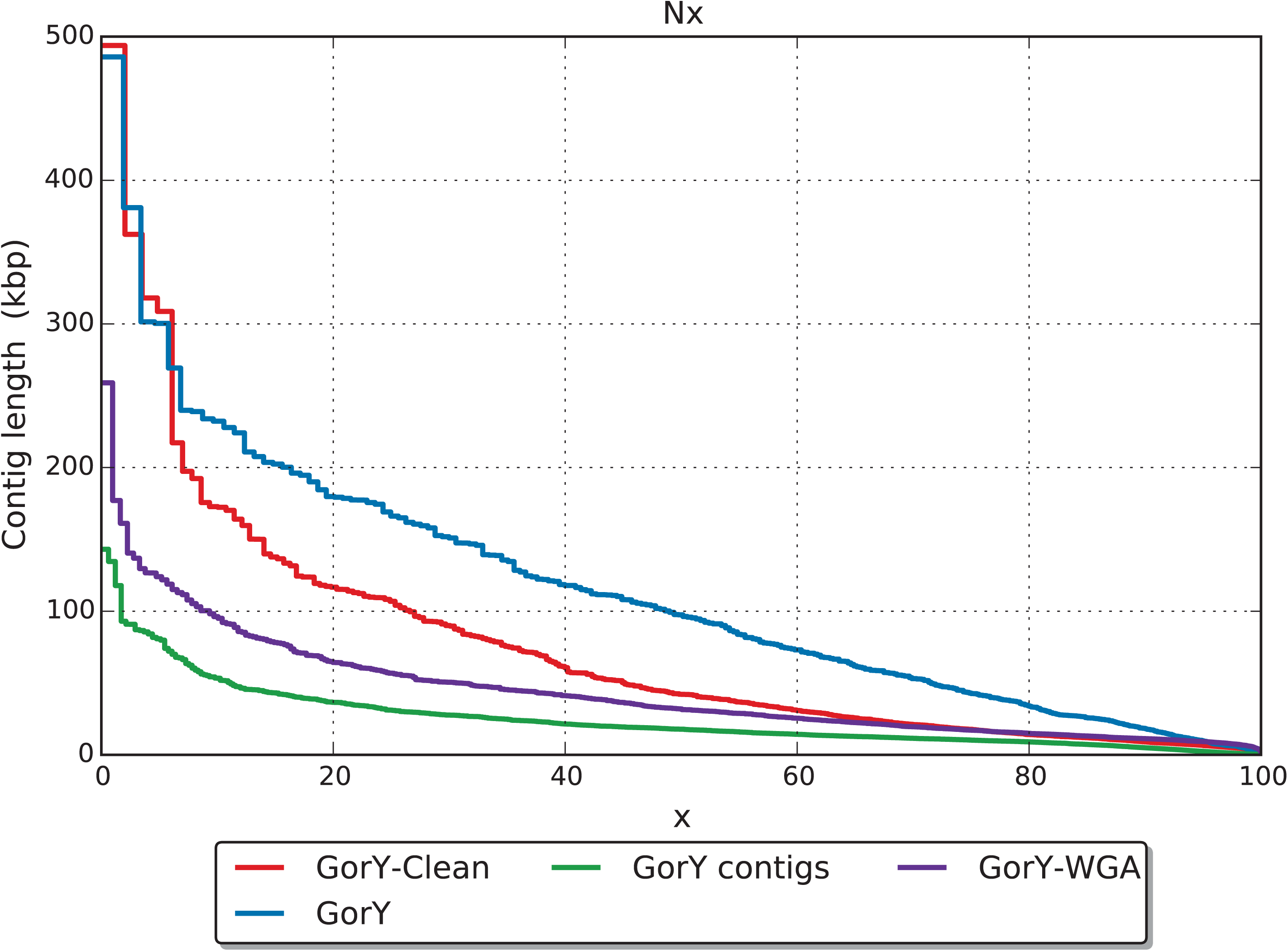
Contig length (y-axis) distribution of the published gorilla Y chromosome (GorY), the contigs underlying this assembly (GorY contigs), the *de novo* assembly based on raw PacBio data set (GorY-WGA) and of the *de novo* assembly of the cleaned reads (GorY-Clean). The x-axis shows the fraction of the assembly (e.g. the N20, N50, etcetera).

**Table 4:** Assembly statistics for the published human and gorilla Y chromosome assemblies and the new assemblies.

|  | Assembly size (Mbp) | Ns (Mbp) | Non-Ns (Mbp) | No. sequences | N50 (Kbp) | Longest seq. (Kbp) |
| --- | --- | --- | --- | --- | --- | --- |
| HumY | 57.2 | 30.4 | 26.8 | 1 |  | 57227 |
| GorY, contigs | 23.0 | 0 | 23.0 | 3001 | 18 | 143 |
| GorY, scaffolds | 25.4 | 2.4 | 23.0 | 697 | 98 | 486 |
| GorY-WGA, contigs | 26.5 | 0 | 26.5 | 1128 | 32 | 256 |
| GorY-Clean, contigs | 24.3 | 0 | 24.3 | 1062 | 42 | 494 |

The accuracy of the newly constructed GorY-Clean contigs become more apparent when looking at the read mapping statistics (Table 5). To calculate these, only contigs are used as these contain sequence information: reads will not map to Ns in the scaffolds. The gorilla Illumina HiSeq reads map better to the *human* reference genome (HumY) than to the original GorY assembly (in terms of genome coverage) and overall mapping accuracy is highest for both newly created assemblies. The GorY-Clean assembly is better covered by the read data than the GorY-WGA assembly is, regardless of whether corrected or non-corrected reads are used for evaluation. These results indicate that, apparently, currently available assemblers (in our case Canu) are better in handling chimeric reads than previous software and that the newly created assemblies (GorY-WGA and GorY-Clean) are more accurate than GorY.

**Table 5:** Mapping of HiSeq, PacBio WGA and PacBio cleaned reads on the human Y chromosome (HumY), the gorilla Y chromosome (GorY) and the newly created gorilla Y assemblies (GorY-WGA, GorY-Clean). The read coverage is the average number of reads that a nucleotide has aligned to it.

| Assembly | Length<br>(Mbp) | HiSeq |  |  | GorY-WGA |  |  | GorY-Clean |  |  |
| --- | --- | --- | --- | --- | --- | --- | --- | --- | --- | --- |
|  |  | Genome coverage |  | Read cov. | Genome coverage |  | Read cov. | Genome coverage |  | Read cov. |
|  |  | (Mbp) | (%) |  | (Mbp) | (%) |  | (Mbp) | (%) |  |
| HumY | 26.8 | 22.3 | 83 | 1897 | 18.2 | 68 | 58.87 | 18.5 | 69 | 74.84 |
| GorY | 23.0 | 18.3 | 80 | 1169 | 21.1 | 92 | 71.33 | 21.1 | 92 | 99.15 |
| GorY-WGA | 26.5 | 24.9 | 94 | 1353 | 26.5 | 100 | 73.15 | 26.5 | 100 | 97.08 |
| GorY-Clean | 24.3 | 22.4 | 92 | 1586 | 24.3 | 100 | 83.21 | 24.3 | 100 | 109.67 |

The average coverage when using the raw reads increased from 73.15× to 83.21× for the GorY-WGA and GorY-Clean respectively and with the corrected reads from 97.08× to 109.67×. These results show that the *de novo* GorY-Clean assembly fits the read data best and, as seen with the *Arabidopsis thaliana* data, mapping accuracy increases after correcting the palindromic reads.

### Resolving artificial duplications provides a higher coverage of genes on the Gorilla Y chromosome

The gorilla Y chromosome contains 12 single-copy X-degenerate genes (Cortez et al. 2014). To evaluate completeness of these genes in the assemblies, their corresponding transcript sequences were mapped to the GorY and GorY-Clean assemblies using the mRNA aligner GMAP (Wu and Watanabe 2005). The resulting alignments were subsequently used to identify the contigs/scaffolds that harbor these genes. For these 12 genes, the transcript coverage was on average higher in GorY-Clean contigs (84.9%) than in GorY scaffolds (74.9%), while sequence identity was similar (Suppl. Table 1, Suppl. Note 1). Additionally, the complete (exons and introns) sequences of the orthologous genes from the Human genome (GRCh38) were aligned to the contigs/scaffolds harboring these genes in the GorY and GorY-Clean assemblies. A visual inspection of the dotplots (Suppl. Figs. 5-10) identified fewer inversions and duplications in the GorY-Clean contigs than in the GorY scaffolds (Suppl. Table 2). In the alignment of the contigs from the GorY-Clean to the GorY scaffolds containing the same genes (Suppl. Figs. 11-16), no inverted duplications were detected in the GorY-Clean sequences. In contrast, numerous inverted duplications were visible in GorY sequences (Suppl. Table 3). Thus, many inverted duplications were resolved in the assembly generated from the sequencing reads corrected with Pacasus, suggesting that such duplications indeed are an artefact of WGA.

## Discussion

Whole-genome amplification is required for sequencing when a biological sample contains insufficient DNA for direct use in library preparation, but the process creates chimeric fragments. We have developed a new method, Pacasus, to correct long, error-rich reads containing such chimeras, based on high-speed Smith-Waterman alignment. We demonstrated the performance of Pacasus in terms of read mapping accuracy and assembly quality, showing that the loss in read length is clearly offset by the removal of incorrect contiguity information. On the Arabidopsis data, the hybrid assembly improves markedly in quality; and on the gorilla data, a PacBio-only assembly on clean reads is even of higher quality than a hybrid assembly including the original reads.

Pacasus has limited effect on the number of nucleotides in the read set and decreases the average read length by less than 50%. A downside is that inverted repeats present in the genome will be treated as chimera, so that the repeat will be split into its separate elements, if the read does not span the full repeat. However, as shown in this manuscript, not all long reads suffer from chimeras. In most cases there will be sufficient reads long enough to cover the inverted repeat and palindrome sequences that are not split by Pacasus. It should be noted that Y chromosomes naturally possess non-artificial palindromes (Skaletsky et al. 2003) and further analysis should be performed to evaluate how processing reads with Pacasus affects the resolution of such palindromes. The gorilla Y chromosome, known to contain palindrome sequences which are also found in the published assembly (Tomaszkiewicz et al. 2016), is assembled after processing with Pacasus, showing that at least some of the inverted repeats with biological origin are still in the long reads.

Flow-sorted chromosomal DNA is usually contaminated with DNA from other chromosomes. Also with the gorilla sample, the original estimate is that approximately a third of the reads originate from the Y chromosome (Tomaszkiewicz et al. 2016). This is supported by our results, with 1,742,887 PacBio reads out of 4,546,488 reads mapping to the GorY-Clean assembly (38%). Consequently, some of the assembled contigs are not part of the gorilla Y chromosome but are from other chromosomes. Further analyses need to be performed to verify which contigs indeed originate from the Y chromosome. One suggestion is to look at read coverage: high coverage could point to Y chromosome sequences (see Suppl. Figure 4d). The next step to further improve the quality of the assembly could be to scaffold the contigs using the RNA-Seq data from the previous study (Tomaszkiewicz et al. 2016) with for example SSPACE (Boetzer et al. 2011) and polish the final assembly with the HiSeq paired-end data using Pilon (Walker et al. 2014).

On the Arabidopsis data, the hybrid assembly improves markedly in quality; and on the gorilla data, a PacBio-only assembly on cleaned reads is even of higher quality than a hybrid assembly including the original reads. The differences between the GorY-Clean and GorY-WGA assemblies are not as large as in the *A. thaliana* case. The underlying reason for this is the much lower number of detectable palindromes in the gorilla read set: 11.8% of the reads contain palindromes, compared to 45.8% of the reads in the *A. thaliana* set. Correcting the relatively low number of reads containing palindromes in the gorilla data set already gave an improvement in assembly statistics, which indicates that the impact of these incorrect reads on the assembly quality is high. We expect that longer reads contain more palindromes, as indicated by the differences in average lengths before and after correcting in both examples.

A possible application not discussed in this paper is the detection of a SMRTbell adapter that is missed by the PacBio software pipeline, producing a raw read with the same structure as created with WGA. These incorrect reads, although perhaps present in low numbers, can have an impact on quality of the de novo assembly. When a non-WGA PacBio dataset with high genome coverage produces a fragmented assembly, it is worthwhile to run Pacasus on this dataset to correct the palindrome sequences created due to the missed SMRTbell adapter

The detection of the palindrome sequences requires a full Smith-Waterman alignment due to the high error rate of the long-read technologies, which takes a considerable amount of compute power. Using high performance software and several GPUs we were able to process one SMRTcell per day, roughly keeping pace with sequencing speed of the PacBio RSII. The throughput of the PacBio Sequel is higher, hence processing these SMRTcells will require more time or compute resources, but we believe the results presented in this manuscript warrant the investment.

To find the location in the read at which it needs to be split, the backtrace part of the Smith-Waterman alignment algorithm needs to be performed (Warris et al. 2015). In the current implementation of the PaSWAS module used for SW, the memory requirements are quadratic in the length of the read. For reads above 100kb this memory requirement may limit the use of Pacasus. Currently the PacBio platforms generate reads below this length, but we expect the Oxford Nanopore platforms to go beyond this limit for at least some the reads in the near future. We will continue to work on Pacasus to decrease the memory requirements of the software to ensure that future output of sequencing platforms can be handled properly.

In summary, Pacasus now allows to analyze PacBio data obtained from low amounts of DNA, making it possible to apply the power of long read technology to, for example, the study of single cells (e.g. in cancer research) and the study of single chromosomes (also in polyploid organisms).

## Methods

### The Pacasus algorithm

To detect chimeras created during WGA, raw PacBio reads are aligned to their reverse-complement sequence with Smith-Waterman (Figure 1) using pyPaSWAS (Warris and Timal 2016). The parameters used for alignment are: gap score, -3; match score, 3; mismatch score, -4. For filtering, the parameters are: filter factor, 0.01; query coverage, 0.01; query identity, 0.01; relative score, 0.01; and base score, 1.0. When no overlap is found in the alignment, the read is left intact and stored in the output file; otherwise the read is split at the center of alignment (see Figure 1(d)). Both resulting fragments are again processed as if they were original reads, to allow the detection of nested palindrome sequences, until no overlap is detected anymore. If a fragment becomes shorter than a minimum length (default 50 bp) it is discarded. Note that the nucleotide information in the reads is neither removed nor changed. Pacasus is implemented in Python 2.7 and, besides pyPaSWAS (version >= 2.0), depends on Biopython (Cock et al. 2009) (version >= 1.67), numpy (http://numpy.org) (version >= 1.8.0) and scipy (Jones et al. 2001) (version >= 0.12.0).

### Data for *Arabidopsis* evaluation

DNA was isolated from two *Arabidopsis thaliana* plants, labeled “Ath-WGA1” and “Ath-WGA2”, and amplified using the REPLI-g Mini Kit (QIAGEN Benelux BV, Venlo, The Netherlands). The *Arabidopsis thaliana* are in-house samples based on low-input plant materials. Both samples were sequenced on both an Illumina HiSeq2000 sequencer and a PacBio RSII sequencer. A third DNA sample was used to generate PacBio RSII data without WGA (“Ath-Ctrl”). Table 6 shows the number of reads generated by each platform and for each sample. To evaluate mapping performance, PacBio reads were mapped to the TAIR10 Arabidopsis thaliana *Columbia* reference genome (http://www.arabidopsis.org) using BLASR version 1.3.1.124201 (Chaisson and Tesler 2012), and alignments with identity < 80% were filtered out by a Python script. Mapping reports were generated in CLCBio version 8.0.2 (http://www.clcbio.com).

**Table 6:** Datasets used for the performance analysis of Pacasus.

| Species | Sample | WGA | Illumina HiSeq2000 |  | PacBio RSII |  |
| --- | --- | --- | --- | --- | --- | --- |
|  |  |  | reads | length | Reads | avg. Length |
| *Arabidopsis thaliana* | Ath-WGA1 | yes | 31233196 | 100 | 462138 | 9326 |
|  | Ath-WGA2 | yes | 43810780 | 100 | 447364 | 8544 |
|  | Ath-Ctrl | no |  |  | 940162 | 5680 |
| *Gorilla gorilla* | GorY-WGA | yes | 279601852 | 150 | 3596236 | 5468 |

PacBio-only *de novo* assemblies of the *A. thaliana* genome were created using Canu version 1.3 (Koren et al. 2017). Hybrid assemblies, combining the HiSeq2000 and RSII data, were created with DBG2OLC (released in 2016) (Ye et al. 2014). DBG2OLC requires as input a HiSeq-only assembly; which we created using Sparse (released in 2015) (Ye et al. 2012), as recommended on the website by the authors of DGB2OLC, based on the HiSeq data from the WGA samples in all cases. The assembly was finalized with the PacBio reads.

We combined Ath-WGA1 and Ath-WGA2 into a single set, Ath-WGA, and created assemblies combining the HiSeq contigs with the raw Ath-WGA reads, with corrected (or ‘clean’) Ath-WGA reads (Ath-Clean) and with Ath-Ctrl reads. To evaluate quality, assemblies were compared to the reference genome using QUAST version 4.3 (Gurevich et al. 2013).

### Data for the gorilla Y chromosome evaluation

PacBio RSII data of a flow-sorted and amplified gorilla Y chromosome, GorY-WGA (Table 1), was downloaded from the NCBI Short Read Archive (SRA SRX1161235). The previously published assembly of the gorilla Y chromosome and the publicly available data from the flow-sorted, whole-genome amplified and de-branched gorilla Y chromosome, GorY (Tomaszkiewicz et al. 2016), were downloaded as well (GCA_001484535.2). Canu version 1.3 was used for the assembly of the PacBio reads. For comparison, the human chromosome Y assembly (NC_000024.10), HumY, was downloaded. QUAST (Gurevich et al. 2013) was used for assembly comparison and statistics. PacBio reads were mapped to HumY, GorY and the new assemblies using BLASR (>80% identity and >80% read coverage); Illumina HiSeq 2500 PE reads (SRA SRR2176191) were mapped using CLCBio version 8.0.2. Statistics for all mapping results were calculated in CLCBio. For calculating the contig length distribution of the GorY assembly, scaffolds were broken up and N’s were removed. The gorilla X-degenerate gene transcripts were retrieved from a previous study (Cortez et al. 2014). GMAP version 2017-03-17 (Wu and Watanabe 2005) was used to align the transcripts to the assemblies.

## Data access

The software is available at https://github.com/swarris/Pacasus.

The project is available at the European Nucleotide Archive under accession number PRJEB21791. The *Arabidopsis thaliana* HiSeq read sets are available under accessions ERX2095148 and ERX2095149. The PacBio data sets are available under accessions ERX2095150 and ERX2095151. The Gorilla Y chromosome assembly has been assigned accession number GCA_900199665.

## Acknowledgments

This work was carried out on the Dutch national e-infrastructure with the support of SURF Cooperative. The authors acknowledge support from the National Science Foundation (awards DBI-ABI 0965596 to K.D.M.; DBI-1356529, IIS-1453527, IIS-1421908, and CCF-1439057 to P.M.), Eberly College of Sciences and Institute for CyberScience at Penn State, the Penn State Clinical and Translational Sciences Institute, and the Pennsylvania Department of Health, using Tobacco Settlement Funds (the Department specifically disclaims responsibility for any analyses, interpretations or conclusions). RV was supported by the National Institutes of Health (NIH)-PSU funded Computation, Bioinformatics and Statistics (CBIOS) Predoctoral Training Program (1T32GM102057-0A1). We would like to thank Jan-Peter Nap (Hanze University of Applied Sciences, Groningen) for his comments and suggestions on the text.

## Author contributions

SW implemented the Pacasus tool and wrote the draft manuscript together with and supervised by DdR. ES was responsible for setting up the ATH experiments, including the whole-genome amplification of the DNA and identifying the palindrome formation after WGA. TH and BtLH sequenced the ATH samples using the PacBio and Illumina HiSeq platform respectively. SW and HvdG performed the *de novo* assemblies on the ATH data set. SW, ES, HG and GSP were involved in validating the results after correction of the palindrome sequences and in checking the quality of the ATH assemblies. SW created the PacBio-only assemblies of the gorilla Y chromosome. RV, PM and KM did the comparison to the existing gorilla Y chromosome assembly. All authors read and approved the manuscript.

## References

Berlin K, Koren S, Chin C-S, Drake JP, Landolin JM, Phillippy AM, Fostier J, Myers E, Berling K, Koren S, et al. 2015. Assembling large genomes with single-molecule sequencing and locality-sensitive hashing. Nat Biotechnol 33: 623–630.

Boetzer M, Henkel C V, Jansen HJ, Butler D, Pirovano W. 2011. Scaffolding pre-assembled contigs using SSPACE. Bioinformatics 27: 578–579.

Buermans HPJ, den Dunnen JT. 2014. Next generation sequencing technology: Advances and applications. Biochim Biophys Acta - Mol Basis Dis 1842: 1932–1941.

Chaisson MJ, Tesler G. 2012. Mapping single molecule sequencing reads using basic local alignment with successive refinement (BLASR): application and theory. BMC Bioinformatics 13: 238.

Chin C-S, Alexander DH, Marks P, Klammer AA, Drake J, Heiner C, Clum A, Copeland A, Huddleston J, Eichler EE, et al. 2013. Nonhybrid, finished microbial genome assemblies from long-read SMRT sequencing data. Nat Methods 10: 563–9.

Cock PJA, Antao T, Chang JT, Chapman BA, Cox CJ, Dalke A, Friedberg I, Hamelryck T, Kauff F, Wilczynski B, et al. 2009. Biopython: freely available Python tools for computational molecular biology and bioinformatics. Bioinformatics 25: 1422–3.

Cortez D, Marin R, Toledo-Flores D, Froidevaux L, Liechti A, Waters PD, Grutzner F, Kaessmann H. 2014. Origins and functional evolution of Y chromosomes across mammals. Nature 508: 488–493.

Czyz ZT, Kirsch S, Polzer B. 2015. Principles of Whole-Genome Amplification. Methods Mol Biol 1347: 1–14.

Edgar RC, Haas BJ, Clemente JC, Quince C, Knight R. 2011. UCHIME improves sensitivity and speed of chimera detection. Bioinformatics 27: 2194–200.

Gawad C, Koh W, Quake SR. 2016. Single-cell genome sequencing: current state of the science. Nat Rev Genet 17: 175–188.

Gurevich A, Saveliev V, Vyahhi N, Tesler G. 2013. QUAST: quality assessment tool for genome assemblies. Bioinformatics 29: 1072–5.

Hackl T, Hedrich R, Schultz J, Förster F. 2014. proovread: large-scale high-accuracy PacBio correction through iterative short read consensus. Bioinformatics 30: 3004–11.

Human Genome Sequencing Consortium International. 2004. Finishing the euchromatic sequence of the human genome. Nature 431: 931–45.

Jain M, Olsen HE, Paten B, Akeson M. 2016. The Oxford Nanopore MinION: delivery of nanopore sequencing to the genomics community. Genome Biol 17: 239.

Jones E, Oliphant T, Peterson P. 2001. SciPy: Open source scientific tools for Python. http://www.scipy.org.

Koren S, Schatz MC, Walenz BP, Martin J, Howard JT, Ganapathy G, Wang Z, Rasko DA, McCombie WR, Jarvis ED, et al. 2012. Hybrid error correction and de novo assembly of single-molecule sequencing reads. Nat Biotechnol 30: 693–700.

Koren S, Walenz BP, Berlin K, Miller JR, Bergman NH, Phillippy AM. 2017. Canu: scalable and accurate long-read assembly via adaptive k-mer weighting and repeat separation. Genome Res 27: 722–736.

Lasken RS, Stockwell TB. 2007. Mechanism of chimera formation during the Multiple Displacement Amplification reaction. BMC Biotechnol 7: 19.

Lee H, Gurtowski J, Yoo S, Marcus S, McCombie WR, Schatz M. 2014. Error correction and assembly complexity of single molecule sequencing reads. bioRxiv 6395.

Rhoads A, Au KF. 2015. PacBio Sequencing and Its Applications. Genomics Proteomics Bioinformatics 13: 278–289.

Sabina J, Leamon JH. 2015. Bias in Whole Genome Amplification: Causes and Considerations. Methods Mol Biol 1347: 15–41.

Schmuths H, Meister A, Horres R, Bachmann K. 2004. Genome size variation among accessions of Arabidopsis thaliana. Ann Bot 93: 317–21.

Shapiro E, Biezuner T, Linnarsson S. 2013. Single-cell sequencing-based technologies will revolutionize whole-organism science. Nat Rev Genet 14: 618–30.

Skaletsky H, Kuroda-Kawaguchi T, Minx PJ, Cordum HS, Hillier L, Brown LG, Repping S, Pyntikova T, Ali J, Bieri T, et al. 2003. The male-specific region of the human Y chromosome is a mosaic of discrete sequence classes. Nature 423: 825–837.

Tomaszkiewicz M, Rangavittal S, Cechova M, Sanchez RC,Fescemyer HW, Harris R, Ye D, O’Brien PCM, Chikhi R, Ryder OA, et al. 2016. A time- and cost-effective strategy to sequence mammalian Y Chromosomes: an application to the de novo assembly of gorilla Y. Genome Res 26: 530–540.

Walker BJ, Abeel T, Shea T, Priest M, Abouelliel A, Sakthikumar S, Cuomo CA, Zeng Q, Wortman J, Young SK, et al. 2014. Pilon: An Integrated Tool for Comprehensive Microbial Variant Detection and Genome Assembly Improvement ed. J. Wang. PLoS One 9: e112963.

Warris S, Timal R. 2016. pyPaSWAS. https://doi.org/10.5281/zenodo.51155.

Warris S, Yalcin F, Jackson KJL, Nap JP. 2015. Flexible, Fast and Accurate Sequence Alignment Profiling on GPGPU with PaSWAS ed. M. Zhang. PLoS One 10: e0122524.

Wu TD, Watanabe CK. 2005. GMAP: a genomic mapping and alignment program for mRNA and EST sequences. Bioinformatics 21: 1859–75.

Ye C, Hill C, Ruan J, Zhanshan, Ma. 2014. DBG2OLC: Efficient Assembly of Large Genomes Using the Compressed Overlap Graph.

Ye C, Ma ZS, Cannon CH, Pop M, Yu DW. 2012. Exploiting sparseness in de novo genome assembly. BMC Bioinformatics 13 Suppl 6: S1.

Zhang K, Martiny AC, Reppas NB, Barry KW, Malek J, Chisholm SW, Church GM. 2006. Sequencing genomes from single cells by polymerase cloning. Nat Biotechnol 24: 680.

